# Glucocerebrosidase activity and lipid levels are related to protein pathologies in Parkinson’s disease

**DOI:** 10.1101/2022.11.11.516171

**Authors:** Cheryl E. G. Leyns, Alice Prigent, Brenna Beezhold, Lihang Yao, Nathan Hatcher, Peining Tao, John Kang, EunRan Suh, Vivianna M Van Deerlin, John Q. Trojanowski, Virginia M.Y. Lee, Matthew Kennedy, Matthew Fell, Michael X. Henderson

**Affiliations:** Department of Neurodegenerative Science, Van Andel Institute, Grand Rapids, MI 49503; Merck & Co., Inc., 33 Avenue Louis Pasteur, Boston, MA, 02115, United States; Institute on Aging and Center for Neurodegenerative Disease Research, Department of Pathology and Laboratory Medicine, Perelman School of Medicine, University of Pennsylvania, Philadelphia, PA 19104 USA

**Keywords:** α-synuclein, tau, β-amyloid, GCase, glucosylsphingosine, glucosylceramide

## Abstract

Parkinson’s disease (PD) and dementia with Lewy bodies (DLB) are progressive neurodegenerative diseases characterized by the accumulation of misfolded α-synuclein in the form of Lewy pathology. While most cases are sporadic, there are rare genetic mutations that cause disease and more common variants that increase incidence of disease. The most prominent genetic mutations for PD and DLB are in the *GBA1* and *LRRK2* genes. *GBA1* mutations are associated with decreased glucocerebrosidase activity and lysosomal accumulation of its lipid substrates, glucosylceramide and glucosylsphingosine. Previous studies have shown a link between this enzyme and lipids even in sporadic PD. However, it is unclear how the protein pathologies of disease are related to enzyme activity and glycosphingolipid levels. To address this gap in knowledge, we examined quantitative protein pathology, glucocerebrosidase activity and lipid substrates in parallel from 4 regions of 91 brains with no neurological disease, idiopathic, *GBA1*-linked, or *LRRK2*-linked PD and DLB. We find that several biomarkers are altered with respect to mutation and disease severity. We found mild association of glucocerebrosidase activity with disease, but a strong association of glucosylsphingosine with α-synuclein pathology, irrespective of genetic mutation. This association suggests that Lewy pathology precipitates changes in lipid levels related to disease severity.

## INTRODUCTION

Parkinson’s disease (PD) presents clinically as a movement disorder and is diagnosed post-mortem by the loss of dopaminergic neurons in the substantia nigra and the presence of Lewy pathology throughout the brain^1^. Parkinson’s disease progresses to dementia (PDD) in up to 80% of cases^2^ and is pathologically nearly identical to dementia with Lewy bodies (DLB)^3,4,^ suggesting that PD, PDD, and DLB are on a disease spectrum^5,6.^ Lewy pathology is composed of misfolded α-synuclein protein and additional lipids and organelles^7^; these cytoplasmic inclusions are hypothesized to be the result of cells’ inability to clear these toxic proteins. It is not known what precipitates the initial misfolding of α-synuclein, and most PD cases are sporadic, without a known genetic cause^8^. However, common genetic risk variants and rare familial mutations give insight into the development of PD. The most common genetic risk variants for PD lie in the *GBA1* gene^9,10.^

*GBA1* encodes the lysosomal lipid hydrolase, glucocerebrosidase (GCase)^11^. Homozygous mutations in *GBA1* can lead to the lysosomal storage disease, Gaucher disease, due to the accumulation of GCase lipid substrates, glucosylceramide (GlcCer) and glucosylsphingosine (GlcSph), in the lysosome^12^. While heterozygous carriers of the mutations will not develop Gaucher disease, they show an approximately 5-fold elevated risk of developing PD^13^. Remarkably, *GBA1* variant carriers also have an 8-fold elevated risk of developing DLB^14^, making *GBA1* variants the most common risk factor for PD and DLB^13,14,^ and the only genetic risk factor shared across these two α-synucleinopathies.

Variants in *GBA1* elevate risk for both PD and DLB, suggesting that disease risk is directly related to α-synuclein pathology development or clearance. Indeed, neuropathologically, idiopathic and *GBA1*-linked PD and DLB cases present indistinguishably with extensive Lewy pathology^15^. GCase activity is reduced in *GBA1*-PD/PDD/DLB^16^ (**Supplementary Fig. 1**), but to a much lesser extent than in Gaucher disease^12^. The retained GCase activity in *GBA1*-PD seems sufficient to keep lipid substrates largely within normal levels, although elevated GlcCer and GlcSph have been reported in some regions^17-19^. Together, these data suggest that decreased GCase activity and elevated glycosphingolipid levels may exacerbate Lewy pathology.

However, the relationship between GCase and α-synuclein is not unidirectional. Work in cell and animal models has suggested that total α-synuclein levels or misfolded α-synuclein may reduce GCase activity as well^20-22^. Indeed, several studies have found that GCase activity is reduced and glycosphingolipid substrates are elevated in the brains of idiopathic PD patients^16-19,23^ (**Supplementary Fig. 1**). However, these GCase activity and lipid changes have only been observed in certain regions, at certain ages, and have not been found consistently across cohorts^24-27^. Most previous studies have focused on either GCase activity or glycosphingolipid analyses; idiopathic or *GBA1* PD, making it difficult to draw conclusions across studies. Only two studies have examined the relationship of GCase activity to α-synuclein pathology, and both studies found mild negative correlations of pathological α-synuclein with GCase activity^19,23.^

GCase activity has also been explored in the context of other genetic risk factors for PD that impact lysosomal function. Among these is the most common genetic cause of PD, mutations of the *LRRK2* gene^28^. Pathogenic *LRRK2* mutations, including the most prevalent G2019S, increase protein kinase activity and have been associated with altered lysosomal morphology, pH, impaired autophagic flux^29,30,^ and most recently, GCase dysregulation. An initial study on dried blood spots found that GCase activity was increased in *LRRK2* mutation carriers manifesting PD compared to non-carriers^31^. Similarly, elevated GCase activity was identified in PBMCs from LRRK2^G2019S^ carriers manifesting PD relative to healthy controls and subjects with idiopathic PD^32^. However, another study reported reduced GCase activity in patient fibroblasts and iPSC-derived dopaminergic neurons from *LRRK2* mutation carriers^33^, an effect that was reversed with LRRK2 kinase inhibition. Overall, the influence of LRRK2 kinase activity on GCase activity has varied by cell type and methodologies applied, and GCase activity has not been assayed in brain tissue from *LRRK2* mutation carriers.

Together, there is substantial evidence for a role of GCase activity and glycosphingolipid substrates in the etiology of PD and DLB. However, there is still an incomplete understanding of the relationship between genetic status, disease state, GCase activity, lipid levels, and protein pathologies across brain regions. Here, we aimed to gain a systematic understanding of how each of these factors relate by examining neuropathology, GCase activity, and glycosphingolipid levels in parallel across four brain regions of idiopathic PD/PDD/DLB, *GBA1*-PD/PDD/DLB, *LRRK2*-PD/PDD and matched controls. We found that GCase activity was reduced in *GBA1*-PD, but not in idiopathic or *LRRK2*-PD. GlcSph was elevated in *GBA1* and idiopathic cases, especially in individuals with dementia. Importantly, we found that GlcSph was highly correlated with both α-synuclein and tau pathologies, which themselves are highly inter-correlated, suggesting that glycosphingolipid accumulation may occur downstream of protein pathology.

## RESULTS

### Parallel assessment of neuropathology, GCase activity, and lipid levels in idiopathic and genetic PD

We sought to examine the relationships of neuropathology to GCase activity and related lipid levels in genetic and idiopathic α-synucleinopathies. We identified 28 *GBA1* mutation carriers and 7 *LRRK2* mutation carriers with available frozen tissue. To enable the best comparison between idiopathic and genetic cases, we selected 37 idiopathic cases that were matched for age, post-mortem interval (PMI), sex, and disease to the genetic cases. We also identified 19 non-neurologically impaired controls that were matched as closely as possible following the same criterion (**Fig. 1A**). We collected frozen tissue from four regions of each brain—cingulate cortex, frontal cortex, putamen, and cerebellum. Cingulate cortex, frontal cortex, and putamen were selected as regions that each exhibit substantial Lewy pathology, but without extensive neurodegeneration that could impact readouts. The cerebellum was chosen as a comparator region lacking Lewy pathology. Each of these regions has also been examined in previous studies, enabling direct comparisons (**Supplementary Fig. 1**). Tissues were chipped frozen, but fine dissections were done on thawed tissue to collect parallel pieces for histology, GCase activity and lipid analysis by mass spectrometry. For tissue not used for histology, gray matter was carefully resected from white matter to avoid contamination of myelin, which has different sphingolipid content than gray matter^34^.

**Figure 1.**
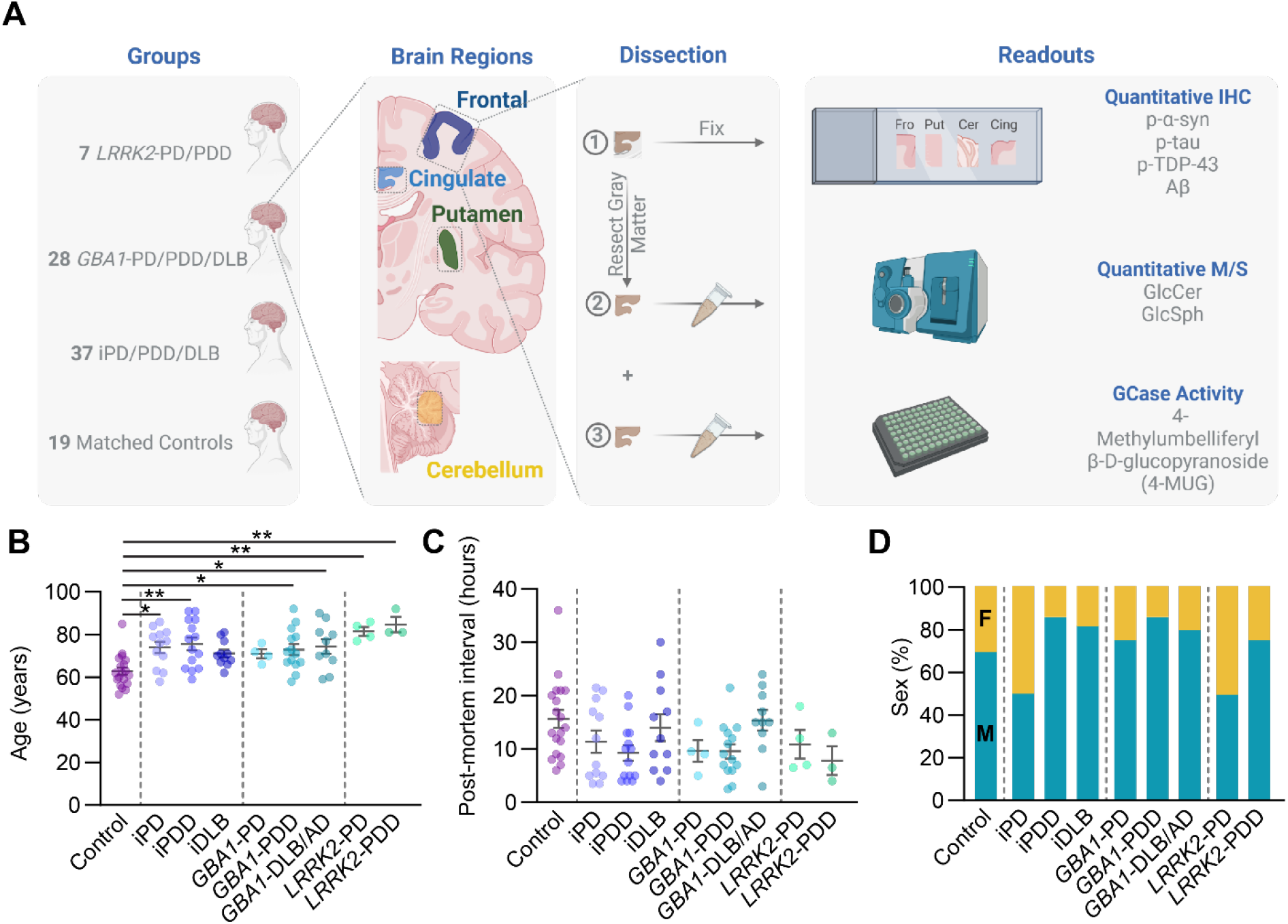
Parallel assessment of neuropathology, GCase activity, and lipid levels in idiopathic and genetic PD. (**A**) Study design. Post-mortem brain tissue was taken from four groups (LRRK2-PD/PDD, GBA-PD/PDD/DLB, idiopathic PD/PDD/DLB and matched controls. Four regions of brain were microdissected, leaving parallel sections for histology, lipid analysis or GCase activity analysis. Samples used for lipid or GCase activity analysis had white matter carefully resected away. (**B**) Ages of subjects at death. (**C**) Post-mortem interval of tissues. (**D**) Percentages of total group represented by each sex. Male (M) are blue-green and female (F) are orange. One-way ANOVA; Tukey’s multiple comparison test. *p<0.05, **p<0.01.

Age for control cases was significantly lower than several of the disease groups (**Fig. 1B**). This is a general feature of control tissue in the brain bank and suitable tissue from control cases of older ages could not be identified. No other major differences were observed between disease cohorts, although *LRRK2* cases were on the older range. Post-mortem interval was well-matched between groups (**Fig. 1C**). The majority of *GBA1* cases were male and sex was well-matched for all cohorts (**Fig. 1D**).

### Quantitative neuropathology of idiopathic and genetic PD

White matter was retained on tissue used for histology to enable proper orientation for NBF, paraffinized, mounted on slides, and stained for the four major aggregating proteins associated with neurodegenerative disease—pS129 α-synuclein, pS202/T205 tau, Aβ, and pS409/410 TDP-43. Gray matter was annotated on each slide (**Fig. 2A**) to enable a direct comparison to GCase activity and lipid levels. Antibodies were selected due to the high signal and low background (**Fig. 2B**) which enabled automated pixel thresholding to quantify area occupied by each of the stains (**Fig. 2C**). Quantification of the percentage of area occupied with pathology spanned several log-fold and enabled pathological comparisons between cohorts (**Fig. 2D**). Control tissues had minimal pathology, other than several cases that had substantial Aβ pathology. Cerebellum also served as an appropriate outgroup, as almost no pathology was observed in this region. Idiopathic and genetic PD/PDD/DLB cases mostly had substantial Lewy pathology in the regions examined, with the exception of a couple *LRRK2*-PD cases, which have been reported to exhibit variable Lewy pathology^35^.

**Figure 2.**
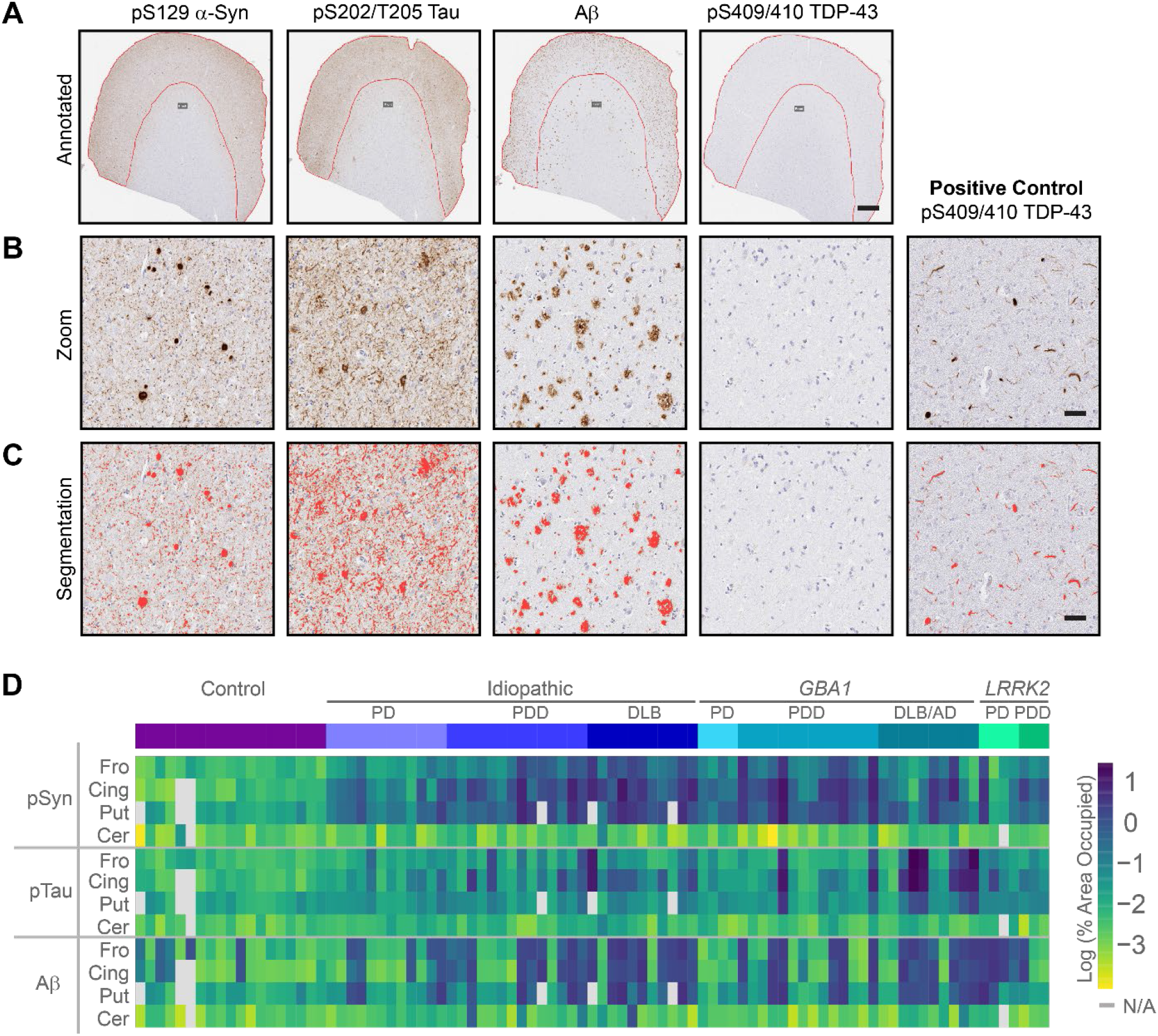
Quantitative neuropathology of idiopathic and genetic PD. (**A**) Representative staining and annotation of gray matter from the cortex of a subject with abundant Lewy, tau, and Aβ pathology. This individual, as for all cases tested, had no apparent TDP-43 pathology. Scale bar = 1 mm. (**B**) A zoomed in image of gray matter showing abundant Lewy bodies, tau tangles, and Aβ plaques. A positive control tissue also shows abundant TDP-43 pathology. Scale bar = 50 μm. (**C**) The same images as in panel B but overlaid with a pixel detection classifier in red at the optimized threshold settings. Scale bar = 50 μm. (**D**) A heatmap of pathology measures from all assayed tissue. Tissues that were not available for assessment are indicated in gray.

To further examine the relationship between pathology and disease cohort, we compared pathology in the cingulate cortex across patient groups. Lewy pathology (pSyn) was elevated in every group except *LRRK2*-PD, compared to control tissues (**Fig. 3A**). Further, in the idiopathic group, there was elevated Lewy pathology in iPDD and iDLB, compared to iPD (**Fig. 3A**). Tau pathology was also elevated in idiopathic and *GBA1* groups (**Fig. 3B**). Aβ exhibited a striking bimodal distribution with an increased prevalence of high Aβ cases in groups with dementia (**Fig. 3C**). We also assessed the relatedness of each pathology type to the other in all examined tissues. pSyn and pTau pathology were highly correlated, with a few notable regions with high pTau and low pSyn, largely from the *GBA1-* DLB/AD group (**Fig. 3D**). The burden of pSyn pathology was also highly correlated with Aβ (**Fig. 3E**), although regions with low pSyn/high Aβ or high pSyn/low Aβ were observed.

**Figure 3.**
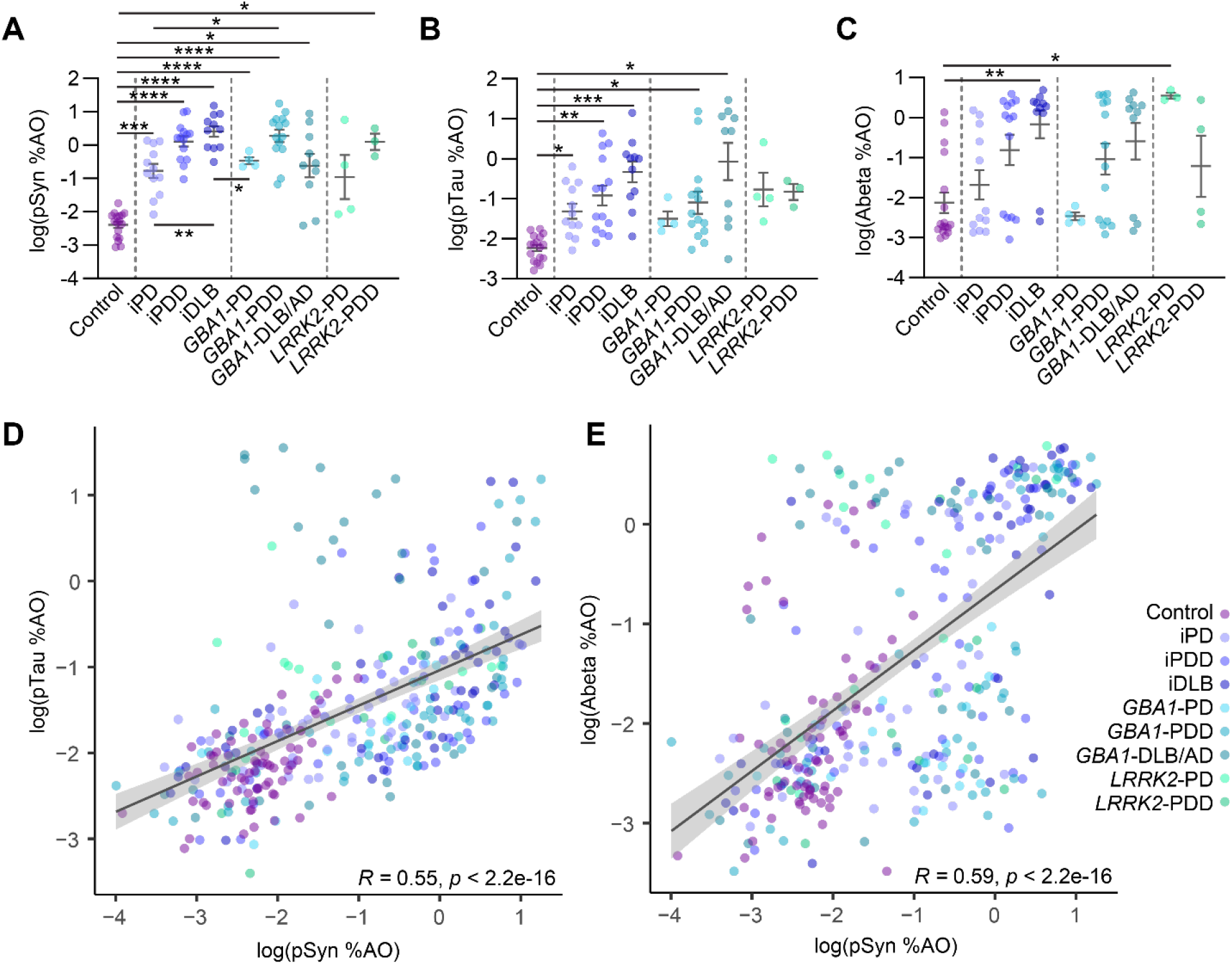
Neuropathological correlations. (**A**) pS129 α-synuclein (pSyn) levels in the cingulate cortex. (**B**) pS202/T205 (AT8, pTau) levels in the cingulate cortex. (**C**) Aβ levels in the cingulate cortex. (**D**) pSyn and pTau levels are highly correlated across all tissues measured. Outliers with high pTau and lower pSyn are largely *GBA1*-DLB/AD cases. (**E**) pSyn and Aβ levels are also correlated across all tissues, but with a bimodal distribution of Aβ. Lines represent linear regression line of best-fit and shaded area is the 95% confidence interval. Panel **A**-**B**: Welch’s ANOVA test; Dunnett’s T3 multiple comparisons test. **C**: One-way ANOVA; Tukey’s multiple comparison test. *p<0.05, **p<0.01, ***p<0.001, ****p<0.0001.

### GCase activity in genetic and idiopathic PD

We next assayed GCase activity in all tissues. GCase activity was determined via the 4-methylumbelliferyl-β-D-glucopyranoside (4-MUG) assay using lysed tissue. GCase activity across patient groups was normalized to control brain tissue levels for ease of interpretation, and when individual regions were analyzed, values were normalized to control brain of only that region. For GCase activity and lipid analyses, two cases were removed that had either homozygous (N370S) or compound heterozygous (N370S, R463C) *GBA1* genotypes due to the dramatically different GCase activity and lipid levels for these tissues. We first tested whether GCase activity differed between regions (**Fig. 4A**). There were large differences by region, the highest GCase activity in the cingulate, moderate activity in the putamen and cerebellum and the lowest activity in frontal cortex. GCase activity was significantly lower in *GBA1* cases, irrespective of brain region (**Fig. 4B-4E**). While there were other changes in mean GCase activity levels, there were no significant differences in the idiopathic group, and *LRRK2* cases had no apparent change in GCase activity relative to controls (**Fig. 4B-4E**). We next evaluated the relationship of GCase activity to pSyn pathology. Overall, there was a very slight negative correlation of GCase activity with pSyn levels (**Fig. 4F**). This negative correlation was more easily observed in individual regions other than the cerebellum (**Fig. 4G**), however there was substantial variability in individual residuals.

**Figure 4.**
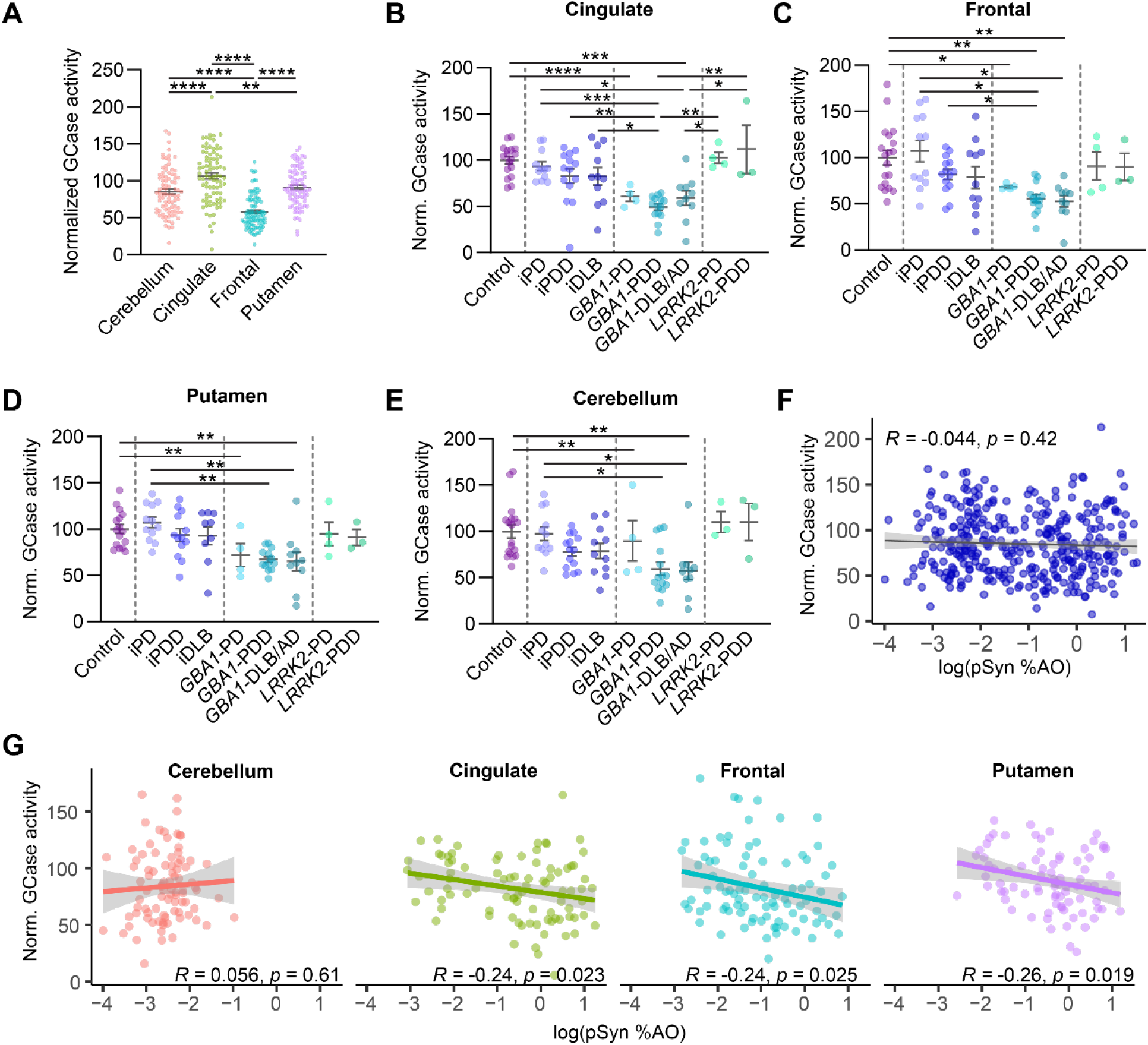
GCase activity in genetic and idiopathic PD. (**A**) GCase activity for all cases, normalized to all control case measures, separated by region. GCase activity is subsequently normalized by region and broken down by neuropathological disease and genetics for each of the four regions: (**B**) cingulate, (**C**) frontal, (**D**) putamen, and (**E**) cerebellum. (**F**) Log normalized pSyn pathology plotted against normalized GCase activity for all samples. (**G**) Log normalized pSyn pathology plotted against normalized GCase activity but normalized and broken down by brain region. Lines represent linear regression line of best-fit and shaded area is the 95% confidence interval. Panel **C**: Welch’s ANOVA test; Dunnett’s T3 multiple comparisons test. **A, B, D, E**: One-way ANOVA; Tukey’s multiple comparison test. *p<0.05, **p<0.01, ***p<0.001, ****p<0.0001.

### GlcCer in genetic and idiopathic PD

While GlcCer is one of the main substrates of GCase, it does not accumulate to a substantial degree in heterozygous *GBA1* mutation carriers, or in idiopathic PD^18,24-26.^ We first sought to determine how GlcCer is distributed across brain regions. We found that GlcCer was significantly lower in the cerebellum than in the cingulate, frontal, or putamen (**Fig. 5A**). We observed no apparent accumulation of GlcCer in any region or disease group compared to controls (**Fig. 5B-5E**). Interestingly, there was an overall positive correlation of GlcCer and pSyn load (**Fig. 5F**), which seemed largely related to a correlation in the cingulate and frontal cortices (**Fig. 5G**). No similar relationship was observed for stereoisomer GalCer (**Supplementary Fig. 2**).

**Figure 5.**
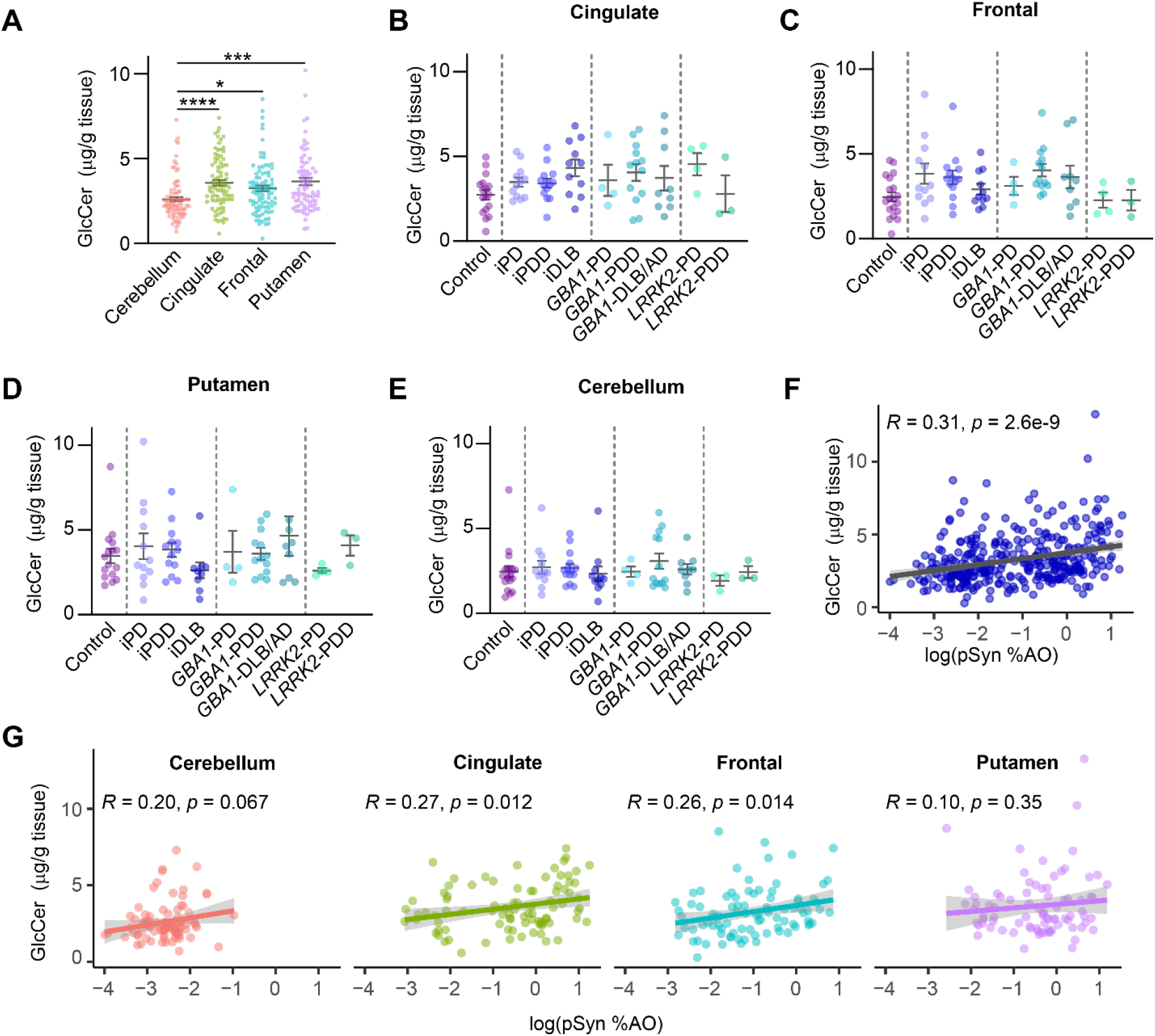
GlcCer in genetic and idiopathic PD. (**A**) GlcCer measures for all cases, separated by brain region. GlcCer levels are subsequently broken down by neuropathological disease and genetics for each of the four regions: (**B**) cingulate, (**C**) frontal, (**D**) putamen, and (**E**) cerebellum. (**F**) Log normalized pSyn pathology plotted against normalized GCase activity for all samples. (**G**) Log normalized pSyn pathology plotted against GlcCer levels but broken down by brain region. Lines represent linear regression line of best-fit and shaded area is the 95% confidence interval. Panel **A**: Welch’s ANOVA test; Dunnett’s T3 multiple comparisons test. **B, C, D, E**: One-way ANOVA; Tukey’s multiple comparison test. *p<0.05, **p<0.01,***p<0.001, ****p<0.0001.

### GlcSph in genetic and idiopathic PD

GlcSph is present at much lower levels than GlcCer, but GlcSph levels have been reported to be increased in *GBA1*-PD and idiopathic PD, albeit to different levels for different regions and dependent on age^17-19^ (**Supplementary Fig. 1**). In contrast to GlcCer, GlcSph is highest in the cerebellum, with moderate levels in the cingulate cortex and putamen, and lowest levels in the frontal cortex (**Fig. 6A**). Individuals with a *GBA1* mutation had higher levels of GlcSph, independent of region (**Fig. 6B-6E**). Within individuals carrying *GBA1* mutations, those with PDD or DLB had GlcSph statistically higher than controls. The *GBA1*-PD group was small and had a lower abundance of the N370S mutation than PDD or DLB/AD groups (**Supplementary Fig. 3A**). The low abundance of mutations other than N370S makes it difficult to make any major conclusions related to specific mutations, and major differences were not observed when GCase activity, GlcSph levels, or pSyn pathology were separated by *GBA1* mutation type (**Supplementary Fig. 3B-3D**) *LRRK2* mutation carriers appeared similar to controls (**Fig. 6B-6E**). In the idiopathic groups, there was a trend for increased GlcSph in the cingulate and frontal cortex associated with increased disease progression. This reached statistical significance in the cingulate cortex, where the iPD group had similar GlcSph levels to controls, but iDLB had significantly higher GlcSph levels than either control or iPD tissue (**Fig. 6B-6E**). This association of GlcSph levels with severity of disease may be related to the burden of pathology in those regions. Considering all samples together, there was no correlation between pSyn pathology and GlcSph (**Fig. 6F**), but this is largely due to the high GlcSph and low pSyn pathology in the cerebellum, as there was a strong correlation between pSyn pathology and GlcSph in the cingulate, frontal, and putamen (**Fig. 6G**). No similar relationship was observed for stereoisomer GalSph (**Supplementary Fig. 4**).

**Figure 6.**
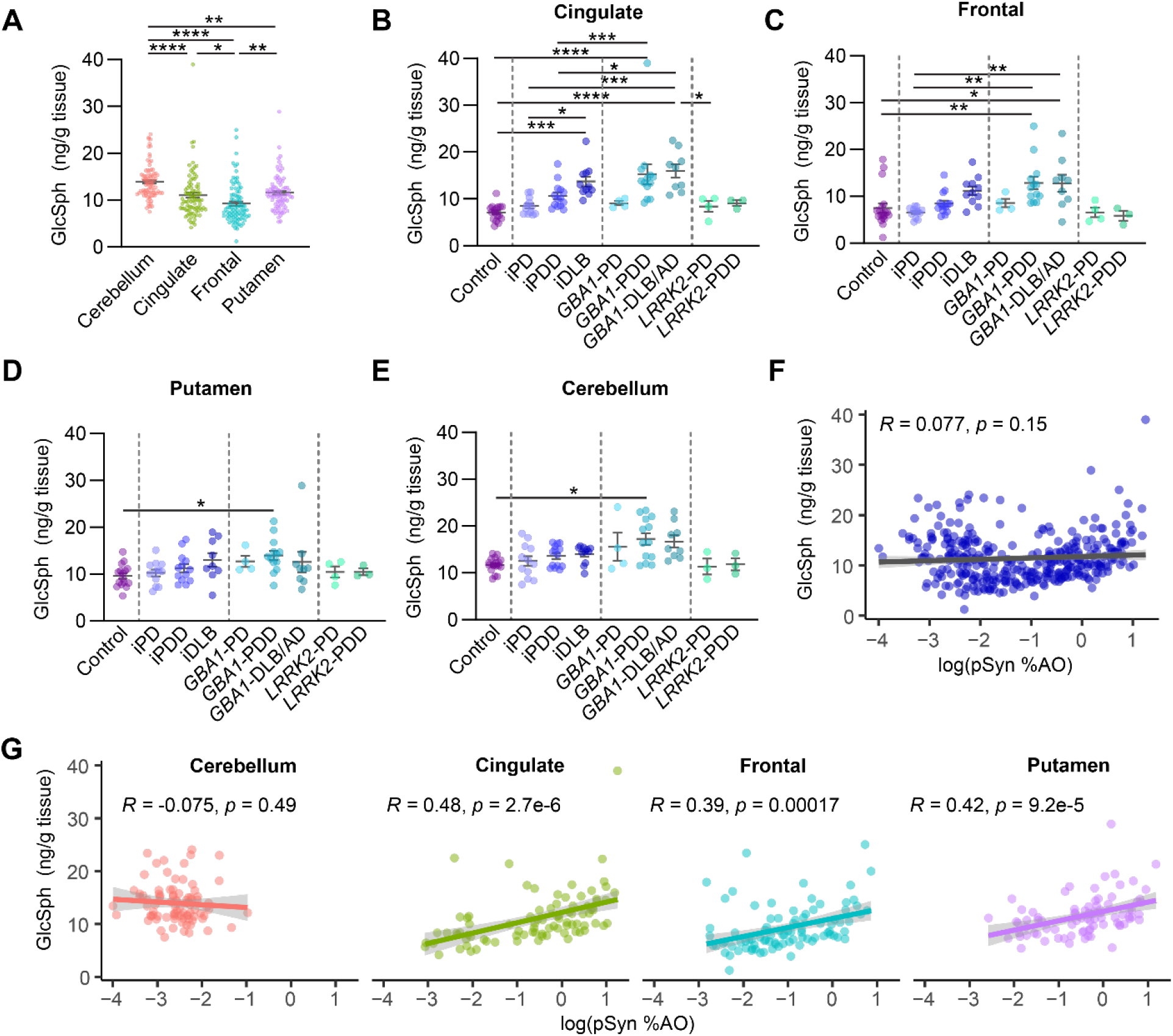
GlcSph in genetic and idiopathic PD. (**A**) GlcSph measures for all cases, separated by region. GlcSph levels are subsequently broken down by neuropathological disease and genetics for each of the four regions: (**B**) cingulate, (**C**) frontal, (**D**) putamen, and (**E**) cerebellum. (**F**) Log normalized pSyn pathology plotted against normalized GCase activity for all samples. (**G**) Log normalized pSyn pathology plotted against GlcSph levels but broken down by brain region. Lines represent linear regression line of best-fit and shaded area is the 95% confidence interval. Panel **E**: Welch’s ANOVA test; Dunnett’s T3 multiple comparisons test. **A, B, C, D**: One-way ANOVA; Tukey’s multiple comparison test. *p<0.05, **p<0.01, ***p<0.001, ****p<0.0001.

### Overall relationships between neuropathology, GCase, and lipids

Given the relationships of protein pathologies with each other and relationships of pathology to GCase activity and lipids, we sought to understand overall relationships across all features measured in this study (**Fig. 7, Supplementary Fig. 5**). Importantly, we also wanted to know if factors such as PMI and age were related to other measures. We found that PMI had no significant correlation with any other measure. GCase activity has previously been reported to decrease with age, with a commensurate increase in lipid substrates^17,18.^

**Figure 7.**
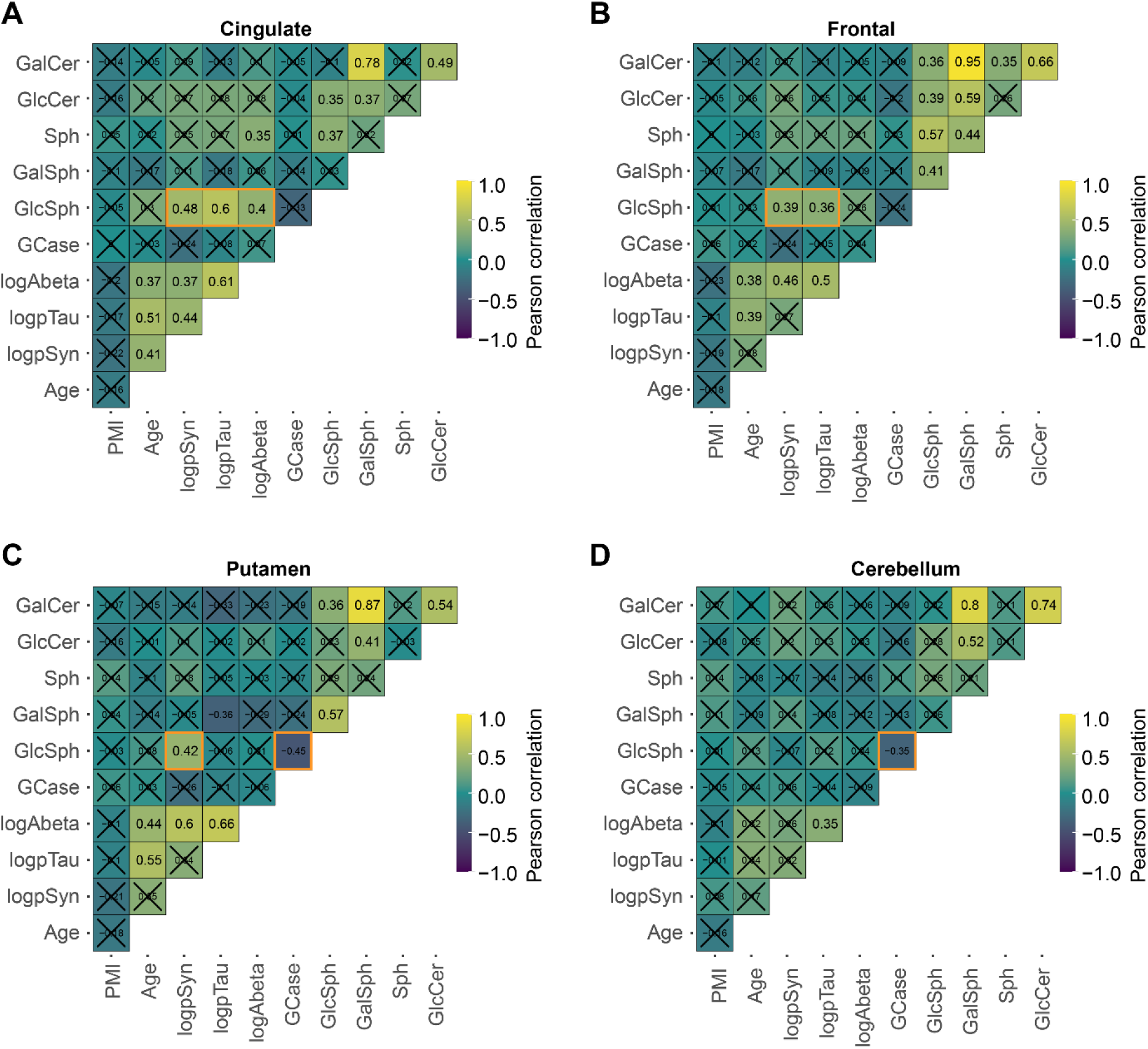
Overall relationships between neuropathology, GCase activity, and lipid levels. Pearson’s correlations between each of the different measure factors, including age, post-mortem interval (PMI), pathology, GCase activity and lipids are plotted here separately for cingulate (**A**), frontal (**B**), putamen (**C**), and cerebellum (**D**). Several of the protein pathologies correlate with each other; several lipid levels also correlate with each other. Of note (highlighted in orange boxes) is the positive correlation between GlcSph and protein pathologies, especially in the cingulate and frontal cortices, and to a lesser extent in the putamen. GCase activity also negatively correlated with GlcSph levels, although this was only significant in putamen and cerebellum. Plots display Pearson’s correlations with Holm’s correction for multiple comparisons. Correlations are noted also by number and those comparisons with p > 0.05 have an “X” over the intersecting box.

We found no significant correlation of age with GCase activity, or glycosphingolipid levels (**Fig. 7, Supplementary Fig. 5**). The age-GCase relationship was primarily observed previously in patients without disease^17^, so we looked more closely in controls at the relationship between age and GCase activity, GlcCer, and GlcSph levels (**Supplementary Fig. 6**). No significant correlation was observed between age and any of these measures.

As previously noted, there were strong correlations of each pathology with each other. Several lipids also showed high correlations with other lipids. The strongest correlation was between GalCer and GalSph. The relationship between GlcCer and GlcSph was notably weaker, only reaching significance in the cingulate and frontal cortex. The strongest correlation with pathology burden was GlcSph levels (**Fig. 7, Supplementary Fig. 5**).

### Variables influencing GlcSph in PD

Due to the apparent associations of GlcSph with pathology, we sought to further delineate the relationships of GlcSph to brain region, GCase activity and pSyn pathology. To examine the contribution of each of these variables in an unbiased manner, we applied a regression decision tree algorithm. A decision tree is a non-parametric supervised learning algorithm used to predict a target variable by learning decision rules from predictor variables. The tree begins with all samples (i.e. root node) for a target variable, GlcSph, and splits on an independent variable that results in most homogeneous sub-nodes (i.e. leaf nodes) of GlcSph values. This partition process is continued recursively. Effectively, each split selects a variable among all variables with the lowest error in predicting GlcSph. The variable importance is then calculated based on the reduction of squared error attributed to each variable at each split and is placed on a scale of 0-100% for each independent variable.

Tree models were built for healthy controls, *GBA1* mutation carriers and idiopathic cases separately. Tree models help capture how variables affected GlcSph regulation. GlcSph levels were associated only by brain regions in healthy, aged controls (**Fig. 8A, Supplementary Fig. 7A**), consistent with the clear regional differences in GlcSph levels (**Fig. 6A**). However, in *GBA*-PD cases, GCase activity was the primary differentiator of GlcSph levels (**Fig. 8A, Supplementary Fig. 7B**), consistent with *GBA1* mutations driving decreased GCase activity and increased glycosphingolipid levels. This was especially true when GCase activity levels were very low. Interestingly, when GCase activity was within a moderate range, GlcSph was co-modulated by pSyn pathology. In idiopathic cases, GlcSph regulation varied regionally. GlcSph was regulated by pSyn pathology, independent of GCase activity, in the frontal cortex only. In the putamen and cingulate cortex, GlcSph was jointly regulated by both GCase activity and pSyn pathology in a complex non-linear fashion (**Fig. 8A, Supplementary Fig. 7C**). As expected, no significant associations were observed between GlcSph and pSyn pathology in the cerebellum in all PD cases. Together, these analyses support the influence of pSyn on GlcSph levels in the presence and absence of *GBA1* mutation and reduced GCase activity.

**Figure 8.**
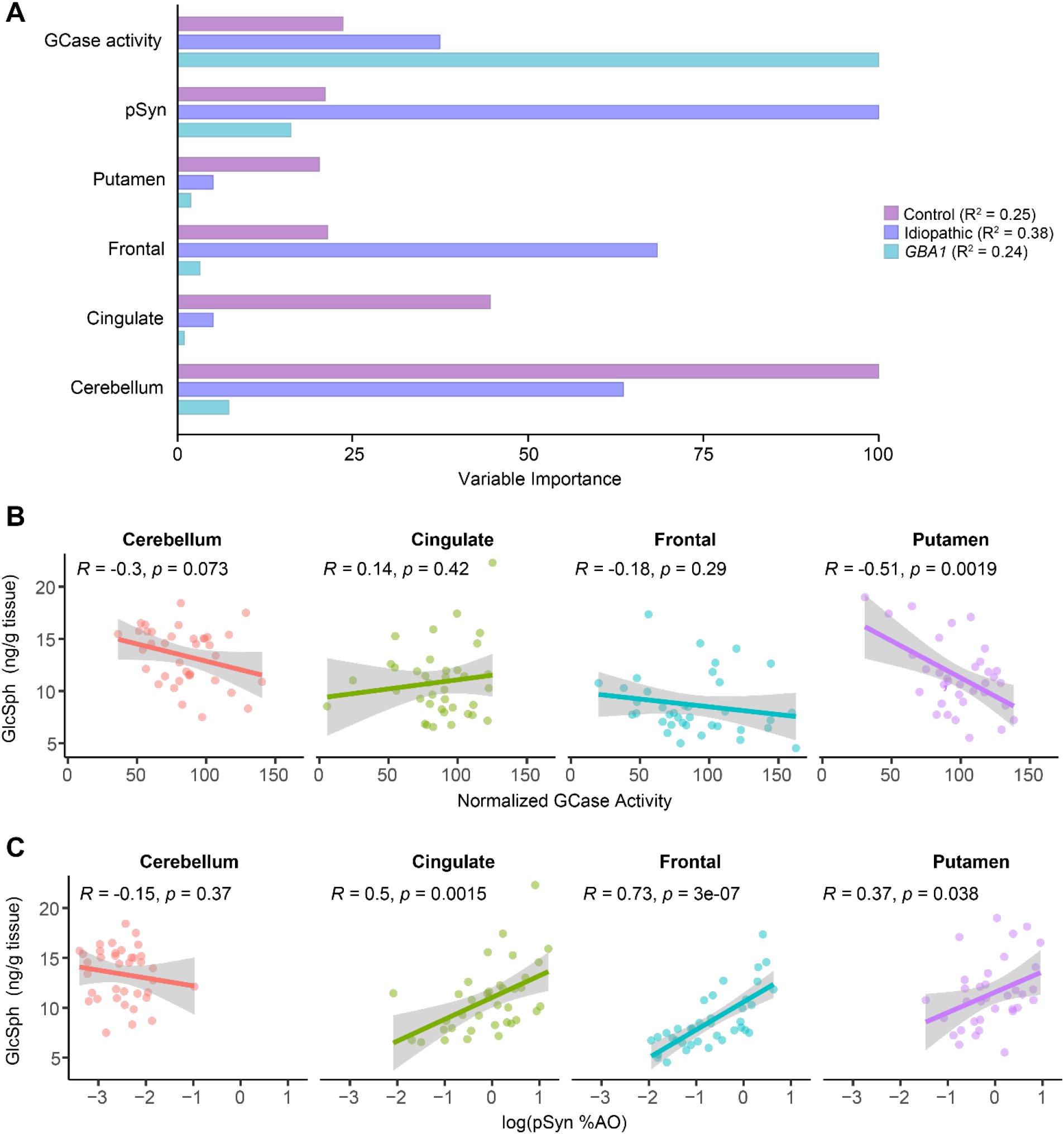
Variables influencing GlcSph levels in PD. (**A**) The importance of variables indicated on the left side were determined and placed within a scale of 0-100 within each group. The R-Squared (R² or the coefficient of determination) is a measure of how well the data fit the regression tree model. Variable importance was determined by calculating the relative influence of each variable contribution to increasing R², i.e., the accuracy of GlcSph prediction. GCase activity (4-MUG) is the most important variable predicting GlcSph in GBA-PD; pSyn is the most important variable predicting GlcSph in iPD; Cerebellum region is the most important variable predicting GlcSph in controls. (**B**) GCase activity for all idiopathic PD, PDD and DLB cases are plotted against GlcSph levels, separated by region. (**C**) Log normalized pSyn pathology for all idiopathic PD, PDD and DLB cases are plotted against GlcSph levels, separated by region. Lines represent linear regression line of best-fit and shaded area is the 95% confidence interval.

The finding that pSyn pathology is a driver of GlcSph levels in idiopathic cases in the tree models prompted us to examine correlations specifically with within idiopathic cases. First, we examined the relationship between GCase activity and GlcSph levels. Outside of the putamen, there is minimal predictivity of GlcSph levels by GCase activity measurements (**Fig. 8B**). However, pSyn pathology showed strong predictivity of GlcSph levels, outside of the cerebellum where there is no pathology (**Fig. 8C**). This is especially prominent in the frontal cortex, as predicted by the tree model. Together, these findings suggest that glycosphingolipids levels are driven by GCase activity in *GBA1* mutation cases, while they are driven by pSyn pathology and GCase activity in idiopathic cases.

## DISCUSSION

*GBA1* variants are the strongest genetic risk factor for developing PD, PDD, or DLB^13,14.^ Yet, most *GBA1*-PD patients retain a wildtype *GBA1* allele and resulting in only modest reductions in GCase activity compared to Gaucher patients DLB^12,16,31.^ Incidence of developing PD for those with a *GBA1* mutation is less than 10%^36,37,^ suggesting that the confluence of factors precipitating disease is likely complex. Multiple studies since 2012 have investigated GCase and its lipid substrates in *GBA1*-PD, and idiopathic PD, to better understand how *GBA1* mutations impact enzyme activity and lipid status, and determine if this enzyme is also impacted in individuals without mutations ^16-19,23-26^ (**Supplementary Fig. 1**). Most studies have individually examined GCase activity, lipid content, idiopathic PD, or *GBA1*-PD. Cohorts have also been differentially segmented by age, mutation, or disease duration, which precludes global summary of these data. However, some major themes have emerged.

GCase activity is reduced in *GBA1*-PD brains, but this is variable and independent of region^16,18.^ This reduction in activity therefore seems related to the mutation itself and less related to disease status. We would hypothesize that healthy *GBA1* mutation carriers would have subtle reductions in GCase activity in the brain though this has not been examined, to our knowledge. Our data are consistent with a reduction in GCase activity across brain regions in *GBA1* mutation carriers, even in regions such as the cerebellum that are largely unaffected by protein pathologies.

Reduced GCase activity has been reported in idiopathic PD^16-18,23,^ but the reported reduction is dependent on brain region and age. If GCase activity is related to Lewy pathology, it would be expected to be reduced only in those regions bearing pathology. However, the cerebellum showed similar reduction of GCase activity in those studies even though cerebellum is typically devoid of α-synuclein pathology. We find no significant reduction of GCase activity in idiopathic PD, although there are certain individuals with low GCase activity, and these tend to be iPDD/DLB groups, consistent with a mild negative correlation between α-synuclein pathology and GCase activity.

Lipid substrates of GCase have been examined in fewer studies, partially owing to the specialized expertise and equipment necessary to isolate GlcCer and GlcSph from their stereoisomers, GalCer and GalSph^25^. In most studies, GlcCer is either unchanged^24-26^ or slightly elevated^18^. Consistent with the literature, we found no consistent change in GlcCer levels in idiopathic or *GBA1* cohorts, suggesting that GlcCer levels are only weakly linked to GCase activity or neuropathology. GlcSph, while it is less abundant than GlcCer, is much more affected in disease. All studies that have examine GlcSph levels in PD brains have found an elevation in GlcSph in some of the regions assayed^17-19^. Our study expanded on these earlier findings, showing that GlcSph levels were elevated in *GBA1* mutation carriers, independent of region, but partially dependent on disease (PD, PDD, DLB). In idiopathic patients, GlcSph showed a strong relationship to disease (PD<PDD<DLB), especially in cortical regions, and a strong correlation with α-synuclein pathology. Further non-parametric analyses identified pSyn pathology burden as the sole regulator of GlcSph levels in the frontal cortex. In the cingulate cortex and putamen, GlcSph was jointly regulated by pSyn and GCase activity (as determined by 4-MUG). Consistent with this analysis, pSyn pathology showed high correlations with GlcSph levels in idiopathic cases in all regions with pathology, while GCase activity only showed high correlations with GlcSph levels in the putamen.

Finally, our study extended the examination of GCase activity and lipid substrate levels to PD subjects carrying *LRRK2* mutations. These tissues were included due to the previous literature showing that LRRK2 kinase activity may be related to GCase activity^31-33,38.^ Despite this compelling literature, we found that the *LRRK2* group had similar GCase activity and lipid levels to control brains, suggesting that *LRRK2* mutations are not driving disrupted GCase activity in the tissues examined. It should be noted that both LRRK2 and GCase are highly expressed in peripheral tissues and may have a stronger interaction there^39^. In addition, only 7 *LRRK2* cases were examined, so future work with larger cohorts will be important to test this relationship.

An additional item often reported in the literature which our study hoped to address is the negative feedback loop between GCase activity and pathological α-synuclein accumulation. It can be difficult to resolve where feedback loops begin once they are in place. While the current study cannot show whether a change in GlcSph or α-synuclein pathology came first, it supports a model in which the loop starts with pathological α-synuclein. 90% of *GBA1* mutation carriers never develop PD^36,37.^ Therefore, carrying a *GBA1* mutation does not necessarily precipitate α-synuclein pathology or PD. However, all patients with *GBA1*-PD/PDD/DLB have α-synuclein pathology^15^, suggesting that it is an integral feature of *GBA1*-PD, and that those individuals who have a *GBA1* mutation will be less protected in the event of α-synuclein accumulation. But the data on GCase activity in idiopathic PD has been variable with some studies showing reduction in certain regions^16,18,23,^ and others showing no change^19^. In the current study, GCase activity trended downward in PDD and DLB, but did not reach significance. However, GlcSph showed an increase in idiopathic patients that corresponded with disease severity. Across cohorts, GlcSph showed a strong correlation with α-synuclein pathology. To determine the mechanistic relationship between Lewy pathology, GCase activity, and glycosphingolipid accumulation, it is useful to focus on PD/PDD/DLB without *GBA1* mutations. In these subjects, GCase is a poor predictor of GlcSph levels, especially in cortical regions. α-Synuclein pathology burden shows a much stronger predictivity of GlcSph levels. These data suggest that GCase itself may not be the direct driver of this relationship. Instead, α-synuclein pathology may disrupt GlcSph degradation, or change its distribution. While GlcSph accumulation in PD cases is mild compared to Gaucher disease, it may be sufficient to drive formation of more pathogenic conformations of α-synuclein^40^.

This study has several limitations. The use of post-mortem tissue precludes the ability to study disease longitudinally. While this is a general difficulty with studying the brain, there are substantial efforts underway to measure GCase activity^41^ and lipid levels from blood and cerebrospinal fluid^26,42,^ in the hopes that these more accessible biofluids may reflect changes in the brain^43^. Another limitation is the number of cases. While this is one of the largest cohorts collected to date, future studies with larger numbers of cases may enable further subtype analysis to see if there are specific patients that are more likely to respond positively to GCase-targeted therapies. Another important consideration is the sampling strategy. Sampling methods are not well-reported across the field, but sampling needs to be done carefully to enable comparisons across groups. White matter, for example, has dramatically different lipid content than gray matter^34^, necessitating careful removal of white matter, as possible. White matter was carefully resected in the current study. We also avoided the substantia nigra pars compact due to the likely shift of cell type and phenotype associated with disease. The dramatic degeneration of this region is associated not only with loss of dopaminergic neurons, but also with gliosis^44,45.^ These factors are likely to shift the GCase and lipid content in severely degenerated regions. However, it is also a limitation of this study that we cannot compare more brain regions. A final related limitation of all studies, including our own, is the bulk nature of the data. Different cells likely have different GCase and glycosphingolipid levels, and by taking all these cells indiscriminately, we may miss important cell-specific effects with analysis of the whole tissue. Use of 4-MUG in an acidic lysate is also limited in its ability to specifically capture lysosomal GCase activity. Future studies would benefit from analyses that retain spatial localization of GCase activity and glycosphingolipid levels.

This study provides the first comprehensive assessment of GCase activity, lipid substrates, and neuropathological assessment from adjacent tissues in PD. We examined this in idiopathic, *GBA1*, and *LRRK2* PD/PDD/DLB. One of the advantages of having a large cohort for most groups in this study is that they could be stratified by disease severity (PD/PDD/DLB). When stratified in this way, there are clear differences in the groups, both in terms of neuropathology, as would be expected, but also in GlcSph levels, which accumulate to a greater extent in PDD/DLB than in PD. There are important remaining questions related to how each of these variables interact. While GlcSph accumulates in idiopathic and *GBA-*PD, the amount of accumulation is minimal compared to that seen in individuals with homozygous *GBA1* mutations. Is this accumulation sufficient to keep a negative feedback loop going? Is GlcSph accumulation a biproduct of lysosomal dysfunction? Does GlcSph accumulate to a much higher level, but only in specific cells that bear the burden of disease? Future studies in human tissue and preclinical models will help clarify the association of GCase, lipid levels and Lewy body disease.

## MATERIALS AND METHODS

### Human brain tissue

A total of 90 brains were used in this study, 18 healthy matched controls, 37 idiopathic PD, 28 GBA-PD and 7 LRKK2-PD. All brain tissues were obtained from the Center of Neurodegenerative Disease Research (CNDR) at the University of Pennsylvania. The study protocol was approved by the local ethics committee and informed consent was obtained from next of kin.

Large brain regions were removed directly from each brain while still frozen. Each of the four brain regions was then thawed on wet ice, and carefully prosected to include only the region of interest (cingulate cortex, putamen, frontal cortex, or cerebellum). A fine slice was transferred to 10% neutral buffered formalin (NBF) for overnight fixation. The remainder of the tissue had white matter carefully removed, and approximately 100-200 mg of gray matter allocated into tubes for subsequent GCase activity or lipid analysis.

The fixed tissue was embedded in paraffin after 24 hours for further histological examination.

### Genetics

Genomic DNA was extracted from brain tissues using QIAamp DNA mini kit (Qiagen, Germantown, MD). Mutations and variants in GBA and LRRK2 were identified by targeted next generation sequencing (NGS) on a neurodegenerative disease-focused panel, which includes genes associated with Parkinson’s disease, as previously described^46^ and alignment of sequence reads and variant calling from NGS were assessed by SureCall software (Agilent, Santa Clara, CA). Identified mutations and variants of interest in *GBA1* and *LRRK2* were confirmed by Sanger sequencing or TaqMan assay (ThermoFisher Scientific).

### Immunohistochemistry

After fixation, brains were embedded in paraffin blocks, cut into 6μm sections and mounted on charged glass slides. All slides were de-paraffinized with 2 sequential xylene baths (5 minutes each) and then incubated for 1 minute in a descending series of ethanol baths: 100%, 100%, 95%, 80%, 70%. After a rinse in distilled water, antigen retrieval was performed by using either formic acid for 5 minutes at room temperature or citric acid pH=6 (Vector Laboratories; Cat# H-3300) for 15 minutes at 95 °C. Slides were allowed to cool for 20 minutes at room temperature and washed in running tap water for 10 minutes. To quench the endogenous peroxidase, slides were incubated in 7.5% hydrogen peroxide for 30 minutes at room temperature. Slides were washed for 10 minutes in running tap water, placed for 5 minutes in 0.1 M Tris buffer pH=6 and then blocked for 1 hour at RT in 0.1 M Tris/2% fetal bovine serum (FBS). Slides were incubated in primary antibody in 0.1 M Tris/2% FBS in a humidified chamber overnight at 4°C. The following antibodies were used: rabbit polyclonal anti-pS129 α-synuclein (EP1536Y) (1:20,000; Abcam, Cat# ab51253), mouse monoclonal anti-pSer202/Thr205 tau (AT8) (1:10,000; life technologies, Cat# MN1020), anti-β-Amyloid (1:200,000, NAB228, Center for Neurodegenerative Disease Research, University of Pennsylvania), rabbit monoclonal anti-pS409/410 TDP-43 (1:20,000, Proteintech, Cat# 80007-1RR). Primary antibodies were rinsed off with 0.1 M tris for 5 minutes and then incubated with the appropriate secondary antibody: goat anti-rabbit (1:1000, Vector, Cat# BA1000) or horse anti-mouse (1:1000, Vector, cat# BA2000) biotinylated IgG in 0.1 M tris/2% FBS for 1 hour at room temperature. Slides were rinsed using 0.1 M tris for 5 minutes, then incubated with avidin-biotin solution (Vector, Cat#PK-6100) for 1 hour. Slides were then rinsed for 5 minutes with 0.1 M tris, then developed with ImmPACT DAB peroxidase substrate (Vector, Cat# SK-4105) and counterstained for 15 seconds with Harris Hematoxylin (Fisher, Cat# 67-650-01). Slides were washed in running tap water for 5 minutes, dehydrated in ascending ethanol baths for 1 minute each (70%, 80%, 95%, 100%, 100%) and incubated in 2 sequential xylene baths (5 minutes each). Slides were mounted with coverslip using Cytoseal Mounting Media (Fisher, Cat# 23-244-256). Slides were scanned at 20X magnification using an Aperio ScanScope XT. The digitized images were then used for quantitative pathology.

### Quantitative pathology

All sections, staining, annotation, and quantification were done blinded to disease and genotype. The digitized images were imported into QuPath software, where gray matter was manually annotated for frontal cortex, cingulate cortex, and cerebellum). Putamen contained interspersed white matter tracts that were not removed. Optical density thresholds of 0.35 were set for each protein (α-synuclein, tau, and β-amyloid) immunostaining so only pathological signal was detected. The percentage of positive area occupied was then measured for each stain. For TDP-43, pathology was detected in the positive control tissue, but not in any of the cases, so no quantification was performed. Linear regressions and one-way ANOVA test followed by Dunn’s post hoc were performed in GraphPad Prism 9.

### GCase activity assay

Tissue lysates were prepared in 300μL ice-cold lysis buffer containing 50 mM Tris–HCl, pH 7.4, 1% (by vol) Triton X-100, 10% (by vol) glycerol, 0.15 M NaCl, 1 mM sodium orthovanadate, 50 mM NaF, 10 mM 2-glycerophosphate, 5 mM sodium pyrophosphate, 1 μg/ml microcystin-LR, and complete EDTA-free protease inhibitor cocktail (Roche, 11836170001). The tissue and buffer were placed in a 2mL round bottom tube (Eppendorf) with a steal bead (Biospec Products, 6.35mm, 11079635C) and homogenized for 90 seconds at 30% amplitude in a Qiagen TissueLyser at 40C. Lysates were centrifuged at 20,000 *x* g for 30 minutes and supernatants were collected. Total protein was determined with a Micro BCA Protein Assay per kit protocol (Thermo Scientific, 23235). To determine Gcase activity, lysates were diluted 1:5 (cingulate cortex, cerebellum, putamen) or 1:10 (frontal cortex) into assay buffer containing 0.25% sodium taurocholate (Cayman Chemical, 16215), 0.25% triton-X-100 (Thermo Fisher, A16046.AP), 1mM EDTA (Millipore, 150-38-9) and citric acid sodium phosphate buffer (pH 5.4). Lysate were incubated shaking, in the presence or absence of conduritol B-epoxide (CBE, Cayman Chemical, 15216), at room temperature for 30 minutes. Seventy-five microliters of 1.25mM 4-Methylumbelliferyl β-D-glucopyranoside prepared in 1% BSA (Research Products Inc., A30075) assay buffer was added to 25μL of lysate. Samples were incubated, protected from light, for 60 minutes shaking at 370C. The reaction was stopped by adding 150μL ice cold 1M glycine (pH 12.5). Plates were read (Ex 355, Em 460) on a BioTek Citation5. Two technical replicates were performed for each sample, with and without CBE, and corrected by background subtraction. GCase activity was quantified as [(Sample^DMSO-treated^ – Sample^CBE-treated^) / total protein] and normalized to the control group.

### Tissue glycosphingolipid analyses

Extraction and quantification of tissue glucosylceramides and glucosylsphingosine was performed as previously described^47^. Briefly, frozen brain specimens supplied as resected grey matter isolates were homogenized in MeOH:H2O (1:1) and normalized across samples by weight. Lipid extraction was performed at room temperature by combining 50 μL of homogenate in with an additional 150 μL of MeOH spiked with d3 glucosylceramide d18:/16:0 and 13C6-glucosylsphingosine standards (Matreya, LLC, State College, PA) at 200 ng/mL and 4 ng/mL respectively prior. Solutions were mixed and 200 μL Acetone:MeOH (1:1) added before brief centrifugation and resuspension with 100 μL of H_2_O. Extract solutions were then subjected to centrifugation at 10, 000 x*g*, and supernatants (250 μL × 2) were transferred to new 96-well plates containing 200 μL MeOH:H_2_O (1:1) per well. Lipid analytes were isolated from supernatants and preconcentrated via either C18 solid phase extraction (isolute C18, Biotage AB, Uppsala, Sweden) for glucosylceramides or strong cation exchange (Oasis MCX, Waters Corp. Inc., Milford, MA) for glucosylsphingosine as previously reported (Hamler et al., 2017). Eluates were evaporated to dryness under gentle N_2_ gas and reconstituted in 50 μL DMSO and 200 μL mobile phase B liquid chromatography buffer (see below). Specimens were processed within the same batch run and analyzed by LC-MS/MS overnight.

Multiple reaction monitoring targeted LC-MS/MS quantification of selected glycosphingolipids was performed using a Waters Acquity UPLC (Waters Corp., Inc.) and SCIEX 5500 QTRAP mass spectrometer (Sciex LLC, Framingham, MA) running in positive ion electrospray mode. Separation was performed using a HALO HILIC 2.7 mm column (Advanced Materials Technology, Inc., Wilmington, DE) and 10 min normal phase LC gradient (mobile phase A: 0.1% formic acid in H_2_O; mobile phase B: 95% acetonitrile, 2.5% MeOH, 2.0% H_2_O, 0.5% formic acid and 5 mM ammonium formate). Transitions for selected endogenous glucosylceramide chain length variants were as follows: C16:0 m/z 700.5 > 264.2, C18:0 m/z 728.6 > 264.2, C20:0 m/z 756.6 > 264.2, C22:0 m/z 784.6 > 264.2, C23:0 m/z 798.6 > 264.2, C24:1 m/z 810.6 > 264.2, C24:0 m/z 812.6 > 264.2 and d3 glucosylceramide d18:1/16:0 reference standard m/z transition was 703.5 > 264.2. Linear calibration curves using d5 labeled glucosylceramide d18:1/18:0 standard (Avanti Polar Lipids, Inc., Alabaster, Alabama; m/z 733.6 > 269.4) were used to estimate concentrations of each of the targeted glucosylceramide fatty acid variants; total glucosylceramide values were represented as the sum concentrations of the C16:0 through C24:1 fatty acid isoforms. Glucosylsphingosine was monitored as a single analyte (m/z 462.4 > 282.1), and concentrations were determined using linear calibration curves of glucosylsphingosine and ^13^C_6_-glucosylsphingosine (m/z 467.4 > 282.1) synthetic standards (Matreya). Peak area and curve fit quantification were performed using SciEx MultiQuant software.

### Decision tree analysis

Single trees were built based on CART algorithm^48^ using R language package, rpart^49^. We built three separate trees for healthy controls, GBA1 mutation carriers and idiopathic PD. The tree model performance and variable importance are evaluated via 10-fold cross-validation using ‘train’ and ‘vmrImp’ functions from caret package^50^. R-squared was calculated to estimate the prediction accuracy for each tree. Variable importance is a relative measure of each variable contribution to accuracy of prediction. It was scaled to a maximum value of 100.

### Statistical analysis

GraphPad Prism software version 9.3.1 (GraphPad Software Inc., La Jolla, CA, USA) was used for pair-wise statistical analysis. The data shown in this study are mean ± standard error of the mean (SEM). For comparison of groups, a Brown-Forsythe test was first applied to test if variances were significantly different. If group variances were not different, a one-way ANOVA was applied with Tukey’s multiple comparison test to determine differences between any groups. If group variances were different, Welch’s ANOVA test was applied with Dunnett’s T3 multiple comparisons to determine if there were differences between groups.

Linear regressions and correlation coefficients were all calculated in R (https://www.R-project.org/)^51^. Correlation matrices were generated using the ‘ggcorrmat’ function in the ‘ggstatsplot’ package in R^52^.

## ACKNOWLEDGEMENTS

We would like to thank the patients and families who participated in this research, without whom this study would not have been possible. We would like to thank Terry Schuck and Brian Alfaro with assistance in tissue collection and the Van Andel Institute Pathology and Biorepository Core for tissue sectioning. We also thank members of our labs for discussions related to this manuscript. Research was supported in part by NIH grant R01-AG077573 to M.X.H. Several images were created with BioRender.com.

## AUTHOR CONTRIBUTIONS

C.E.G.L and M.X.H. conceived and designed the experiments. C.E.G.L., A.P., B.B., L.Y., N.H., E.S., P.T., J.K., and M.X.H. performed experiments and analyzed results. C.E.G.L and M.X.H. wrote the manuscript. All authors have reviewed and approved the manuscript.

## DATA AVAILABILITY

The data that support the findings of this study are available from the corresponding author upon reasonable request.

## SUPPLEMENTARY MATERIALS

**Supplementary Figure 1.**
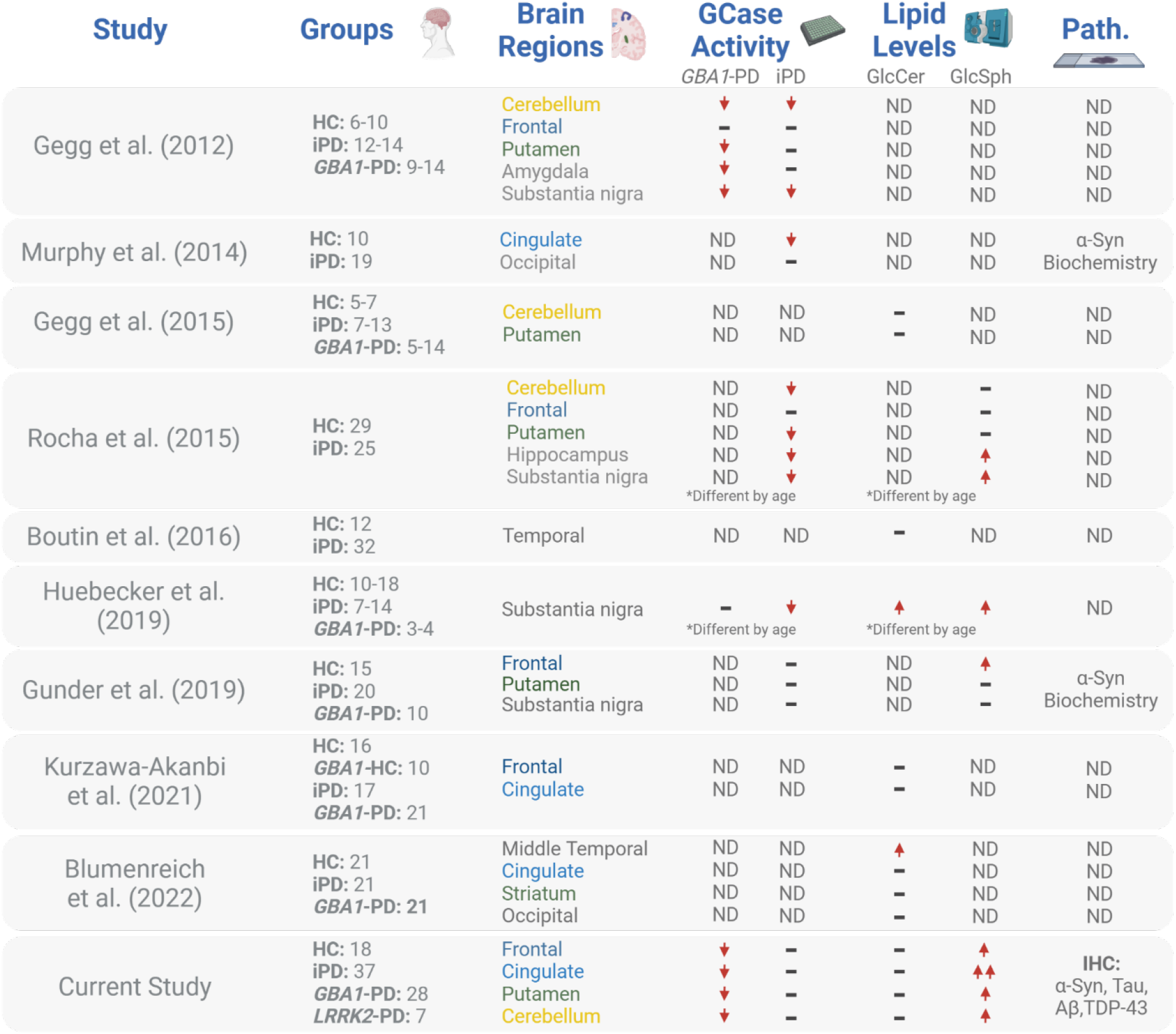
Summary of GCase and lipid studies in human brain. Findings from previous studies that have examined GCase activity, GlcCer or GlcSph lipid levels or pathological proteins in idiopathic or GBA-PD are summarized here. The current study is listed last on the table. ND indicates that this measure was “Not Determined.” Hyphens indicate no significant change in this measure. Down arrows indicate a decrease. Up arrows indicate an increase. The two up arrows in the current study are to indicate the increase of GlcSph in both idiopathic as well as *GBA1*-PD.

**Supplementary Figure 2.**
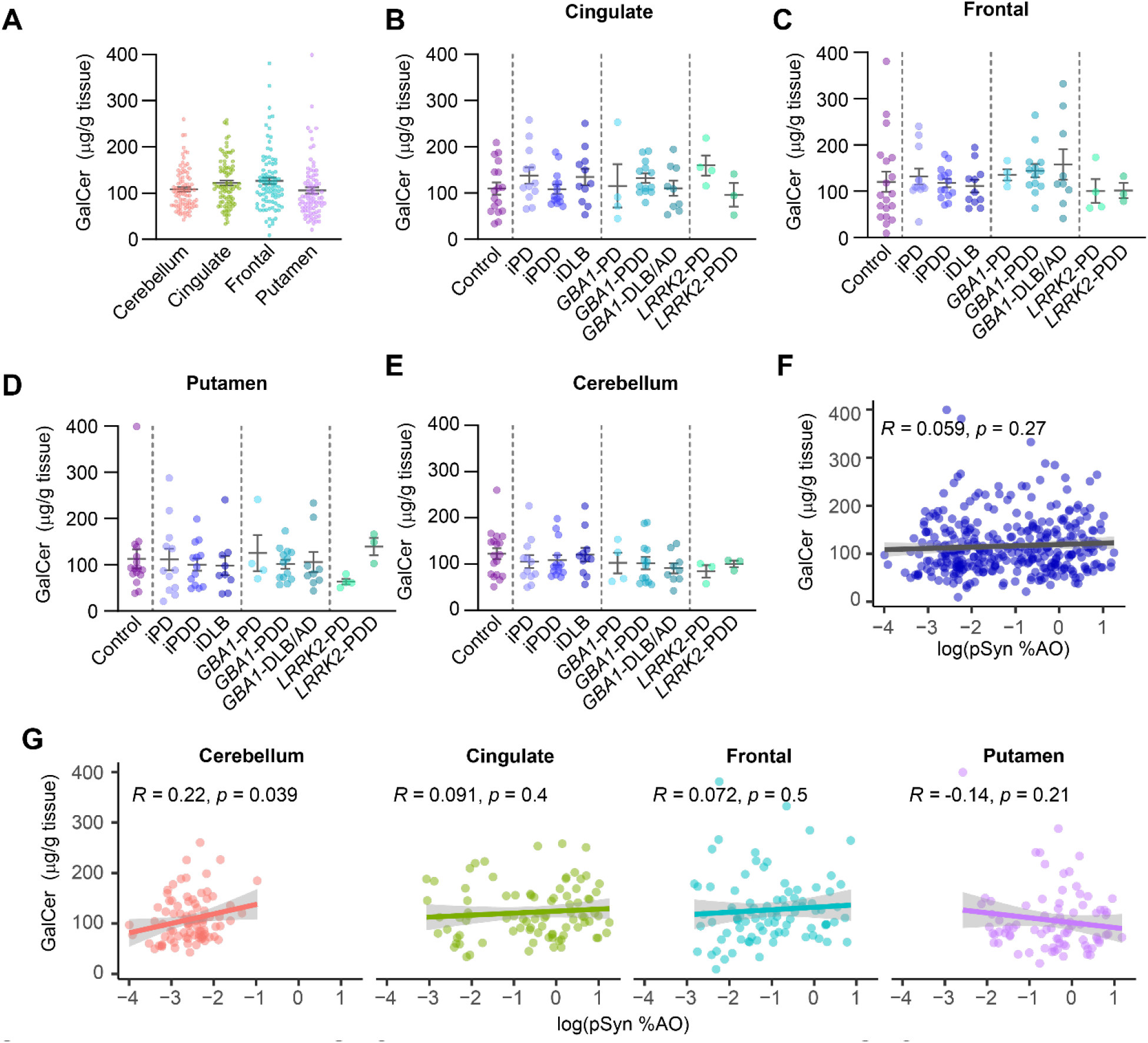
GalCer in genetic and idiopathic PD. (**A**) GalCer measures for all cases, separated by region. GalCer levels are subsequently broken down by neuropathological disease and genetics for each of the four regions: (**B**) cingulate, (**C**) frontal, (**D**) putamen, and (**E**) cerebellum. (**F**) Log normalized pSyn pathology plotted against normalized GCase activity for all samples. (**G**) Log normalized pSyn pathology plotted against GalCer levels but broken down by brain region. Lines represent linear regression line of best-fit and shaded area is the 95% confidence interval. Panels **A, B, C, D, E**: One-way ANOVA; Tukey’s multiple comparison test.

**Supplementary Figure 3.**
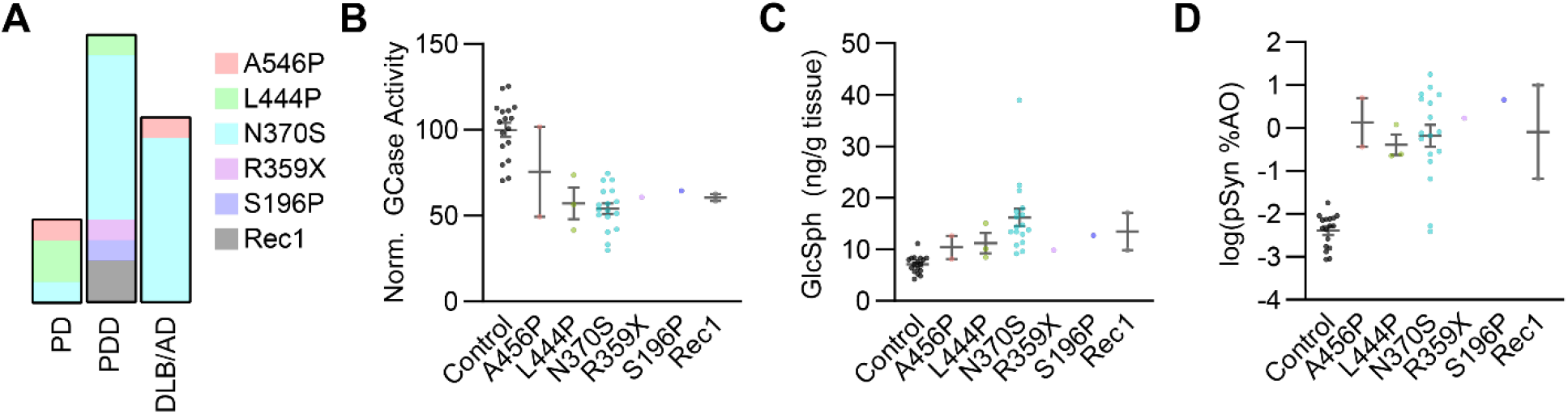
*GBA1* mutation analysis. (**A**) The proportion of all *GBA1* mutation carriers with the noted variant in each disease category is noted. N370S was the predominant mutation overall, and especially for PDD and DLB/AD groups. The PD group was small, but two out of four cases carried L444P. (**B**) Normalized GCase activity in the cingulate cortex for control and *GBA1* mutation carriers. (**C**) GlcSph levels in the cingulate cortex for control and *GBA1* mutation carriers. (**D**) pSyn levels in the cingulate cortex of control and *GBA1* mutation carriers. The number of carriers for individual mutations was insufficient to make statistical comparisons, but no major differences were observed by genotype.

**Supplementary Figure 4.**
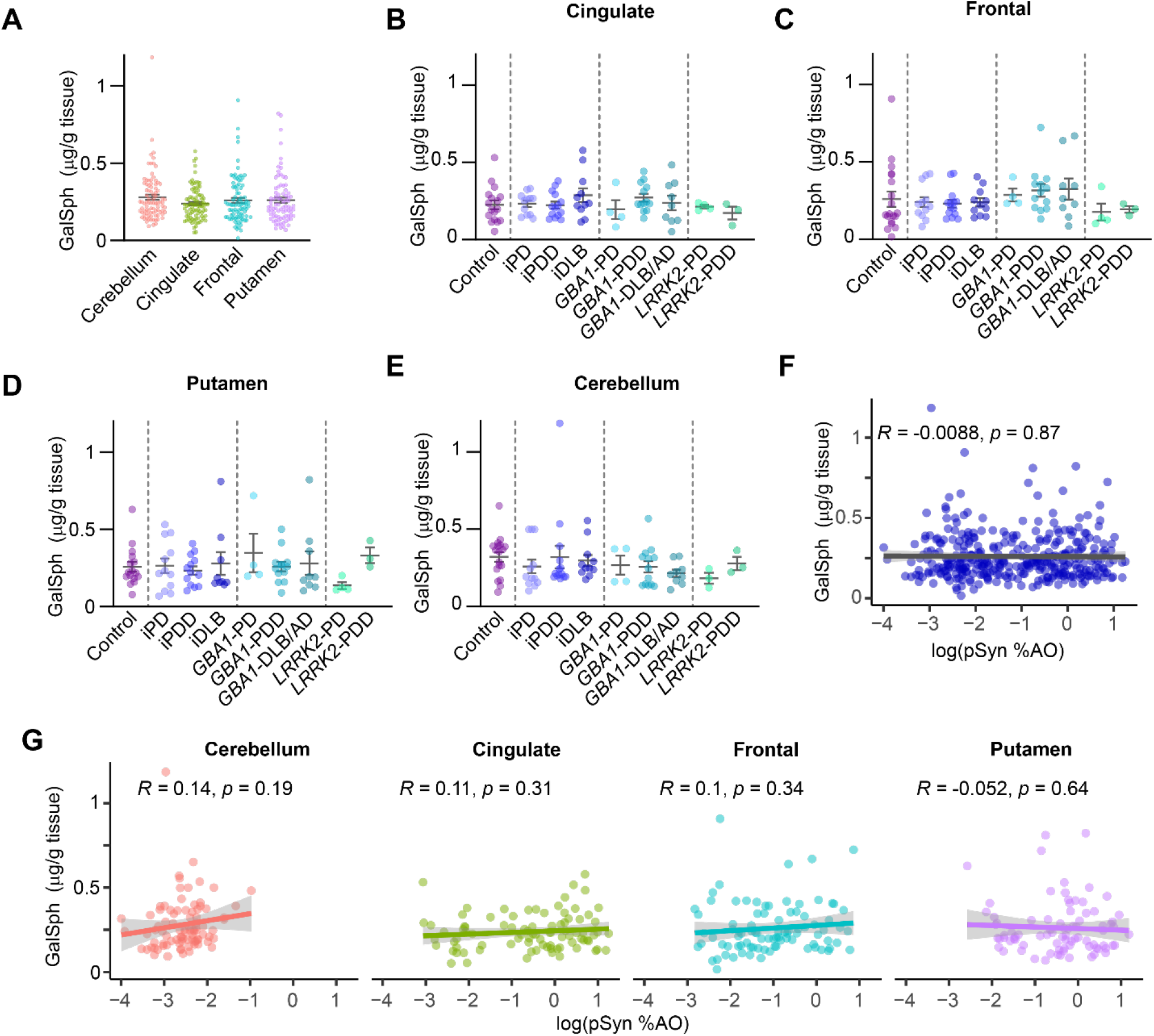
GalSph in genetic and idiopathic PD. (**A**) GalSph measures for all cases, separated by region. GalSph levels are subsequently broken down by neuropathological disease and genetics for each of the four regions: (**B**) cingulate, (**C**) frontal, (**D**) putamen, and (**E**) cerebellum. (**F**) Log normalized pSyn pathology plotted against normalized GCase activity for all samples. (**G**) Log normalized pSyn pathology plotted against GalSph levels but broken down by brain region. Lines represent linear regression line of best-fit and shaded area is the 95% confidence interval. Panels **A, B, C, D, E**: One-way ANOVA; Tukey’s multiple comparison test.

**Supplementary Figure 5.**
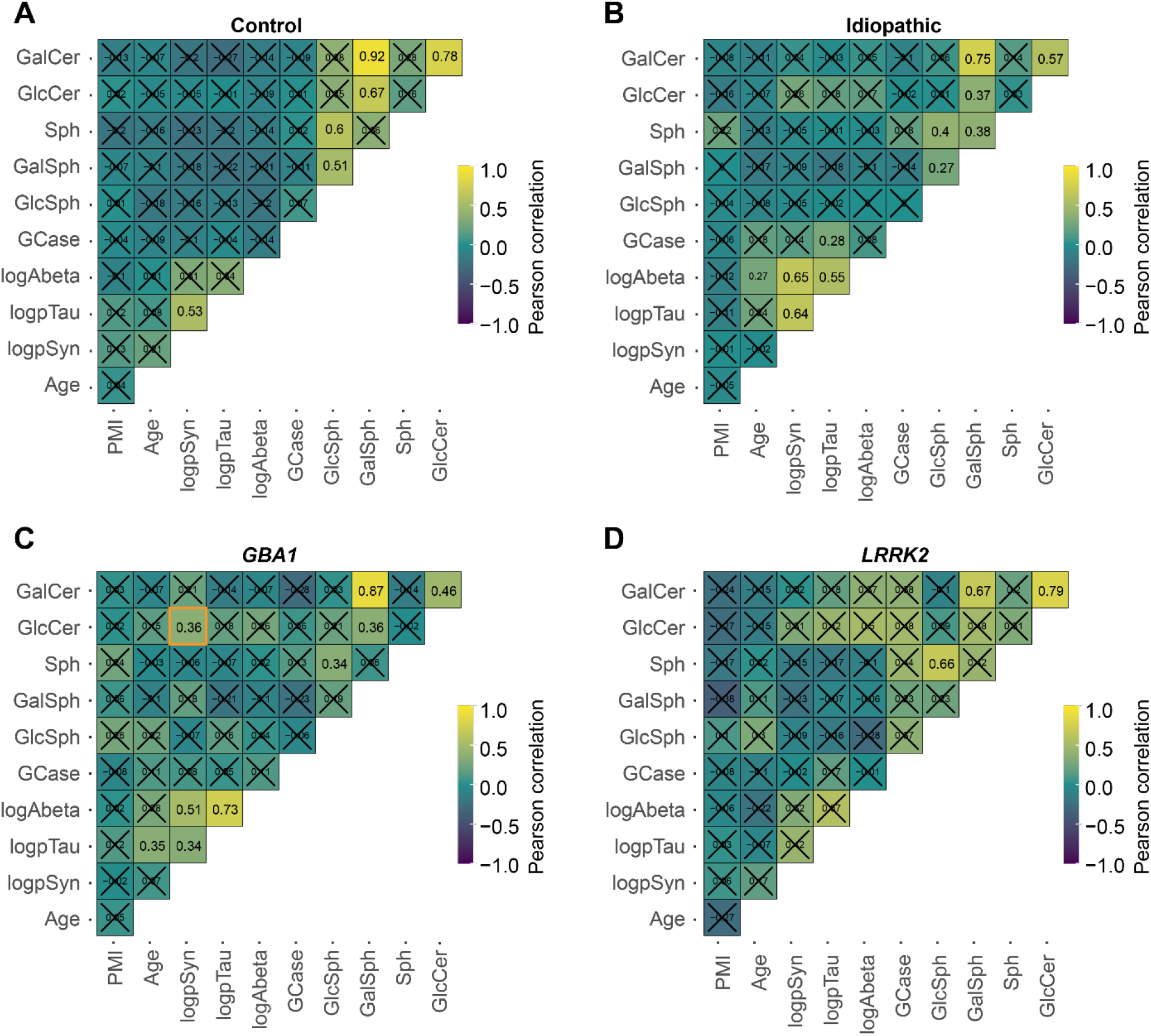
Overall relationships between neuropathology, GCase, and lipids by disease. Pearson’s correlations between each of the different measure factors, including age, post-mortem interval (PMI), pathology, GCase activity and lipids are plotted here separately for control (**A**), idiopathic (**B**), *GBA1* (**C**), and *LRRK2*-linked PD (**D**). Several of the protein pathologies correlate with each other; several lipid levels also correlate with each other. There is minimal correlation with pathologies and lipids within each disease group, with the exception of the correlation of GlcCer with logpSyn in *GBA1*-PD tissue.

**Supplementary Figure 6.**
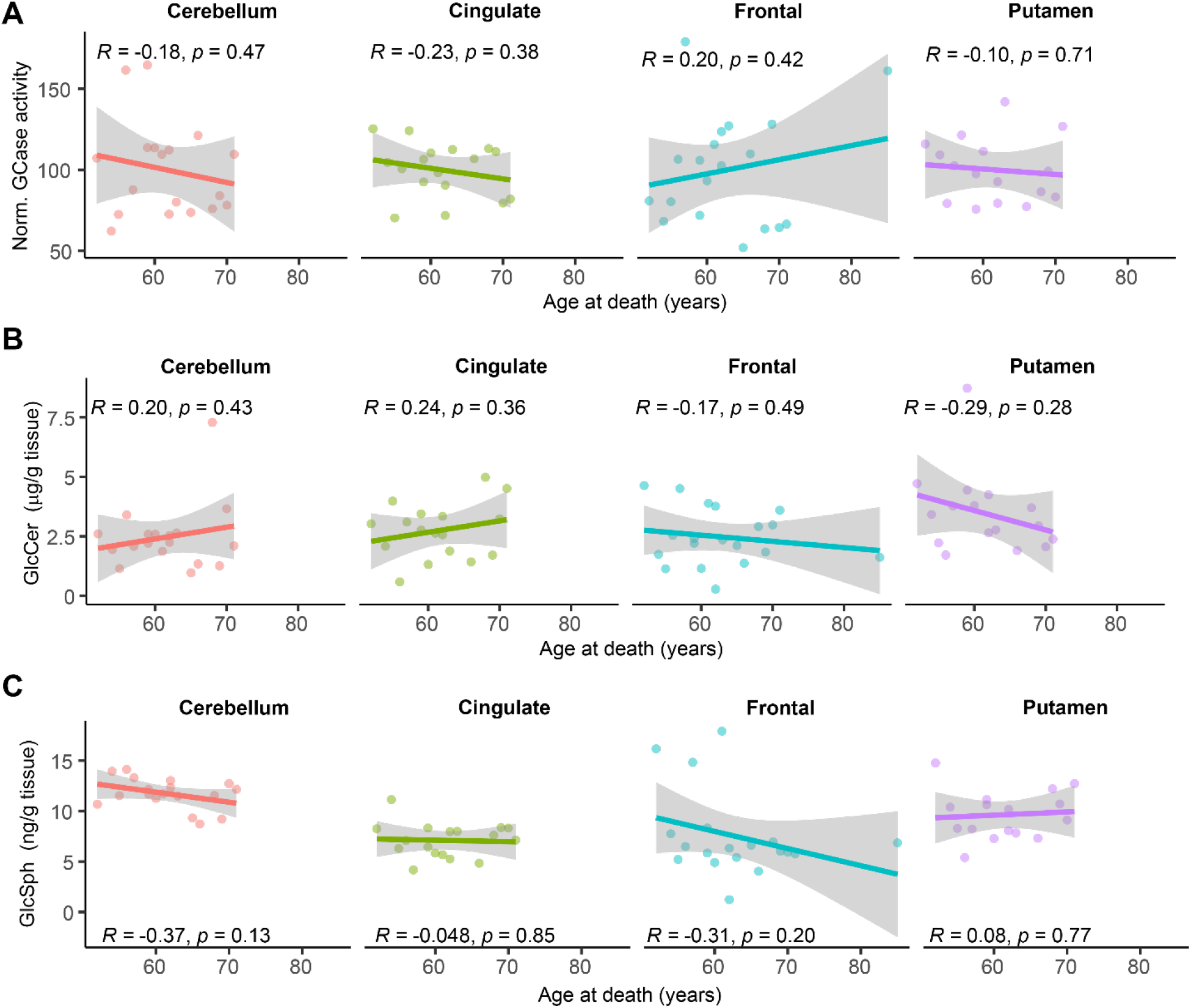
Correlations with age. Age at death for all control subjects for each of the for regions collect was plotted against normalized GCase activity (**A**), GlcCer levels (**B**), or GlcSph levels (**C**). No significant correlations were observed. Lines represent linear regression line of best-fit and shaded area is the 95% confidence interval.

**Supplemental Figure 7.**
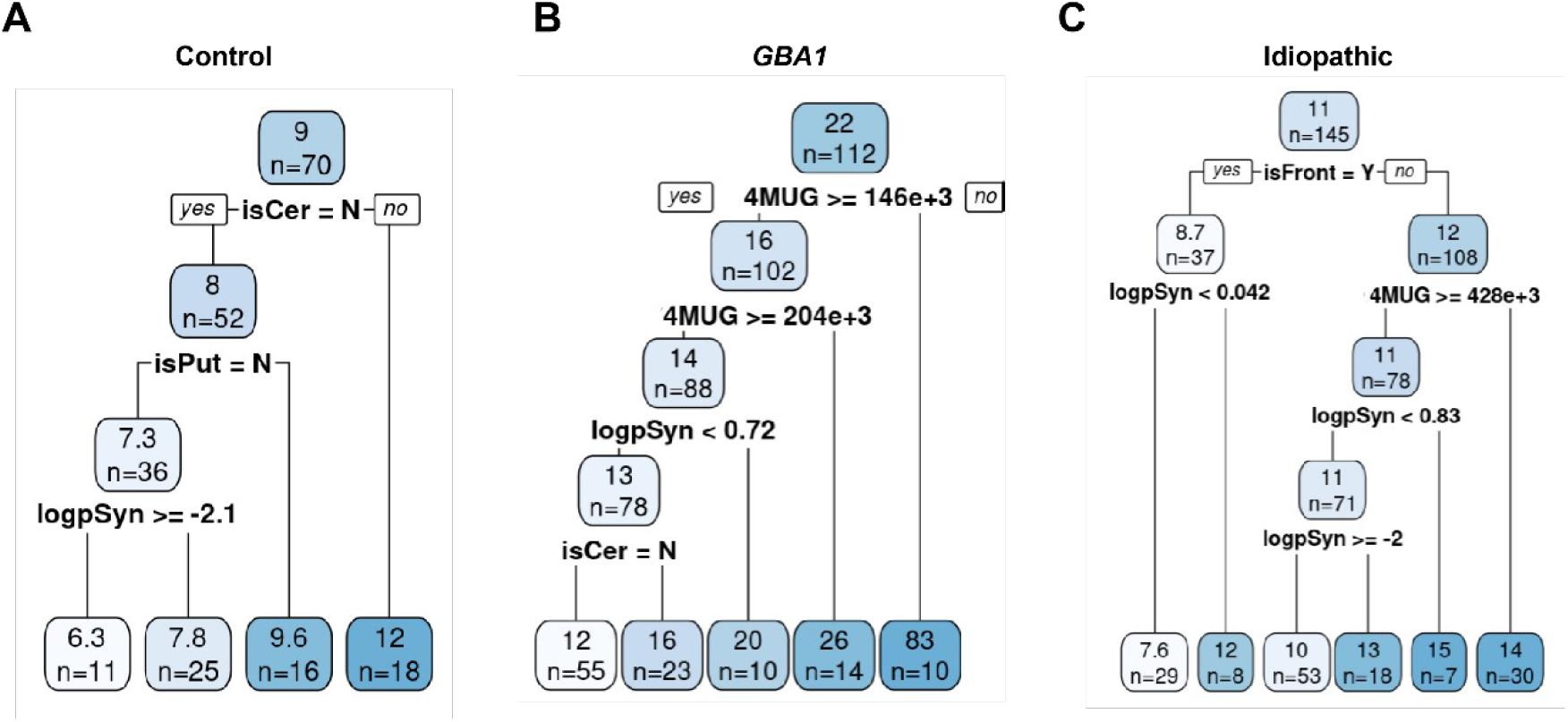
Decision tree analysis of variables influencing GlcSph levels. Three decision trees were generated with (**A**) control, (**B**) *GBA1* and (**C**) idiopathic groups. The root node indicates the average GlcSph value across all samples (sample size indicated below) and each node is then split into left and right sub-nodes. Optimal split variable and associated cut point were shown in each tree up to maximum depth of 5. Tracing the trajectories from root to leaf exhibits insights of the joint interaction pattern of different splitting variables in driving GlcSph. For example, the right path of the decision tree in (**C**) idiopathic group, 4-MUG and pSyn were interacting with one another in non-frontal regions to regulate GlcSph.

